# Manipulating base quality scores enables variant calling from bisulfite sequencing alignments using conventional Bayesian approaches

**DOI:** 10.1101/2021.01.11.425926

**Authors:** Adam Nunn, Christian Otto, Mario Fasold, Peter F. Stadler, David Langenberger

**Author notes:** Authors contributed equally to this manuscript (Adam Nunn, Christian Otto).

## Abstract

Calling germline SNP variants from bisulfite-converted sequencing data poses a challenge for conventional software, which have no inherent capability to dissociate true polymorphisms from artificial mutations induced by the chemical treatment. Nevertheless, SNP data is desirable both for genotyping and to understand the DNA methylome in the context of the genetic background. The confounding effect of bisulfite conversion can be resolved by observing differences in allele counts on a per-strand basis, whereby artificial mutations are reflected by non-complementary base pairs. Herein, we present a computational pre-processing approach for adapting sequence alignment data, thus indirectly enabling downstream analysis in this manner using conventional variant calling software such as GATK or Freebayes. In comparison to specialised tools, the method represents a marked improvement in precision-sensitivity based on high-quality, published benchmark datasets for both human and model plant variants.

## Introduction

DNA methylation is among the most-studied of the molecular mechanisms involved in epigenetics, and has been associated for example with changes in gene expression^1–3^, chromosome interactions^4,5^, and genome stability through the repression of transposable elements^6–8^. It is a base modification most often characterised by the addition of a methyl group to a cytosine nucleotide^9^, to form 5-methylcytosine (5mC) or one of its derivatives e.g. 5-hydroxymethylcytosine (5hmC). Cytosine methylation occurs typically in a CG sequence context in eukaryotes^10^ but is also prevalent in CHG and CHH contexts (where H is any base but G) in plants^11^.

Following the emergence of next-generation sequencing (NGS) technologies, library preparation protocols such as BS-seq^11^ and MethylC-seq^12^ were devised which facilitate the nucleotide-resolution analysis of DNA methylation patterns through the chemical treatment of sample DNA with sodium bisulfite. The treatment catalyses the deamination of unmethylated cytosines to uracil, while methylated cytosines remain unaffected, to produce non-complementary, single-stranded (ss)DNA. As these strands then undergo PCR, uracil pairs with adenosine rather than the original guanosine during replication, which in turn pairs with thymine in the final, amplified product in place of the original cytosine. The resulting paired-end libraries therefore contain four distinct read-types: the forward (FW) and reverse complement (RC) of the bisulfite-converted sequence from the original Watson(+) strand, and the forward and reverse complement of the bisulfite-converted sequence from the original complementary Crick(-) strand. Mapping such reads to the known genome requires specialised software, but when performed successfully can reveal the underlying extent of DNA methylation over each potential 5mC site by considering the proportion of cytosine matches to thymine mismatches. Evidently, any thymine mismatches arising instead as a result of natural mutation are obscured by bisulfite conversion and risk being mistaken as unmethylated cytosines.

Previous attempts to resolve such confounding positions in the genome, to determine both the correct methylation level and reveal single nucleotide polymorphisms (SNPs), have resulted in the development of specialised software such as BISCUIT (https://github.com/huishenlab/biscuit), Bis-SNP^13^, BS-SNPer^14^ and MethylExtract^15^. Each case combines methylation calling and variant calling into a single, concurrent analysis to produce output in a custom variant call format (VCF). No single approach however considers the variant calling itself as a primary, independent outcome. Users looking additionally to leverage SNP data for e.g. genotyping or purposes unrelated to DNA methylation are therefore limited by the scope and rationale behind the development of existing tools. Instead, the present application aims to abstract variant calling as a standalone objective in order to facilitate analysis with conventional software, such as GATK^16^, Freebayes^17^, or Platypus^18^, thereby optimising precision-sensitivity during SNP discovery and allowing users to make the most out of their bisulfite sequencing data for a broader range of purposes.

Under a simple Bayesian framework to variant calling, the conditional probability of observing the true genotype *G* given the variants observed in the sequencing data *D* can be represented for example by equation (1), which formulates the problem as the derivation of a prior estimate of the genotype *P*(*G*) and the likelihood of observing the data *P*(*D*|*G*).

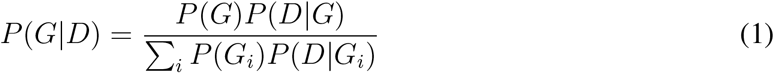

Given that NGS data is seldom error-free, even the simplest model will typically incorporate base quality (BQ) information directly into the Bayesian inference of genotypes as a fundamental scaling factor for the data likelihood estimation. The BQ score itself is a phred-based quality value which denotes on each position the estimated probability that the base caller identified the correct nucleotide during sequencing. In the context of variant calling from bisulfite-treated NGS data, any potential nucleotide conversions present in the resulting sequencing reads can, in principle, be considered analogous to zero-quality base calls. Leveraging this mechanism imposes an indirect strand-specificity on potential variants which cannot otherwise be dissociated from the effect of bisulfite conversion, dictating that they be informed only by opposite-strand alignments where the original, complementary nucleotide is hence unaffected by the treatment.

The method presented herein involves a simple “double-masking” procedure which manipulates specific nucleotides and BQ scores on alignments from bisulfite sequencing libraries (Figure 1), with the formal procedure on individual alignments described in Algorithm 1. It involves two steps which are performed *in silico*. First, specific nucleotides in bisulfite contexts are converted to the corresponding reference base in order to obscure likely methylation sites from consideration as potential variants. This vastly reduces the number of positions that are even considered by the variant caller, where there is no evidence of a SNP beyond the bisulfite treatment. Second, any given nucleotide which may potentially have arisen due to bisulfite conversion is assigned a base quality (BQ) score of 0. This drives the variant caller to make the correct decision in regards to genotype on positions where there is real evidence of a SNP. In paired-end sequencing, the procedure applies in a C>T context on mate 1 alignments to the Watson strand (FW+) and mate 2 alignments to the Crick strand (RC-), whereas mate 1 alignments to the Crick strand (FW-) and mate 2 alignments to the Watson strand (RC+) follow G>A context. Reads obtained from single-end sequencing behave in equivalent manner to mate 1 in paired-end sequencing.

**Figure 1:**
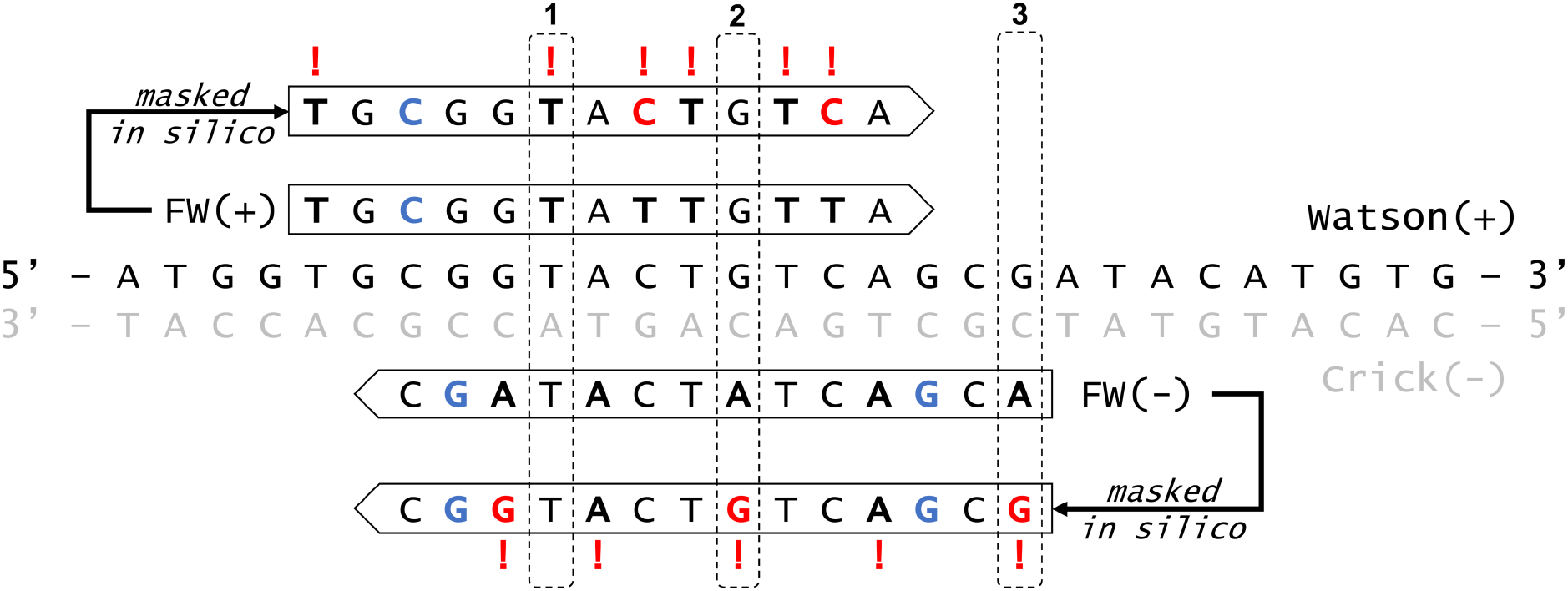
Overview of the double-masking procedure. The central sequence represents the reference genome, with example alignments (+FW and -FW) adjacent to each originating strand. Black, emboldened nucleotides potentially arise from bisulfite treatment. Blue colouring indicates 5mC/5hmC. Red colouring represents *in silico* nucleotide manipulation, and corresponding base quality manipulations are indicated with an exclamation mark. In example **(1)** the variant caller is informed only by the -FW alignment, and in **(2)** only by the +FW alignment. As there is no equivalent Watson(+) alignment in **(3)** it is impossible to determine whether the apparent G>A polymorphism arises from bisulfite or a natural mutation.

In contrast to previous approaches with bisulfite data, the method is applied as a pre-processing step prior to variant calling, thereby facilitating interoperability with conventional, state-of-the-art variant calling software. For validation, SNPs derived from published, experimental whole genome bisulfite sequencing (WGBS) data in human (NA12878) and *Arabidopsis thaliana* (Cvi-0) accessions are compared to high-quality variant standards and high-confidence regions obtained from the NIST Genome in a Bottle initiative^19^ and the 1001 genomes project^20^, respectively. The method presented herein has been implemented as a standalone python script available at https://github.com/bio15anu/revelio, which is intended to be adapted and “plugged-in” to any variant pipeline working with bisulfite data so that the user can choose whichever alignment and variant calling software best suits their purposes. An open-source example of a working pipeline for whole genome data is available at https://github.com/EpiDiverse/SNP, which is itself a branch of the EpiDiverse Toolkit^21^. The software is also implemented by epiGBS2 in the analysis of reduced-representation bisulfite data^22^.

## Results

In benchmark data sets for both test species, precision-sensitivity of the SNPs derived from WGBS data is demonstrably improved following double-masking in comparison to existing methods (Figure 2). Notably, common filtering metrics such as variant quality (QUAL) and genotype quality (GQ) behave as could be expected in conventional sequencing data (Figure 2; Supplementary Figure 1), facilitating in many cases the use of established best-practice criteria for selecting high-confidence calls. Additional comparison of SNPs derived from real WGS data (*A. thaliana*; accession Cvi-0) and equivalent WGBS data, following *in silico* bisulfite conversion (~99%) of sequencing reads, removes the variation caused by differences during sequencing, but not alignment. The resulting ROC-like curves demonstrate a comparable level of sensitivity (i.e. true positives) in both WGS and WGBS data following variant calling with Platypus, Freebayes and GATK3.8 UnifiedGenotyper (Figure 3), however there is a drop in precision driven in each case by an influx of false positives. When *in silico* bisulfite conversion is instead applied directly to the WGS alignments, thus eliminating variation due to the alignment of bisulfite-treated reads, the differences in false positives are reduced for each tool. All software demonstrate an appreciable performance, with GATK3.8 achieving the highest raw number of both true and false positives, followed by Freebayes and then Platypus, for both WGS and WGBS data. The total number of false positives derived from *in silico* WGBS alignments however represent only 1.0%, 3.8% and 4.3% of the total, unfiltered calls for those same tools respectively, when discounting the fraction shared in the equivalent WGS data.

**Figure 2:**
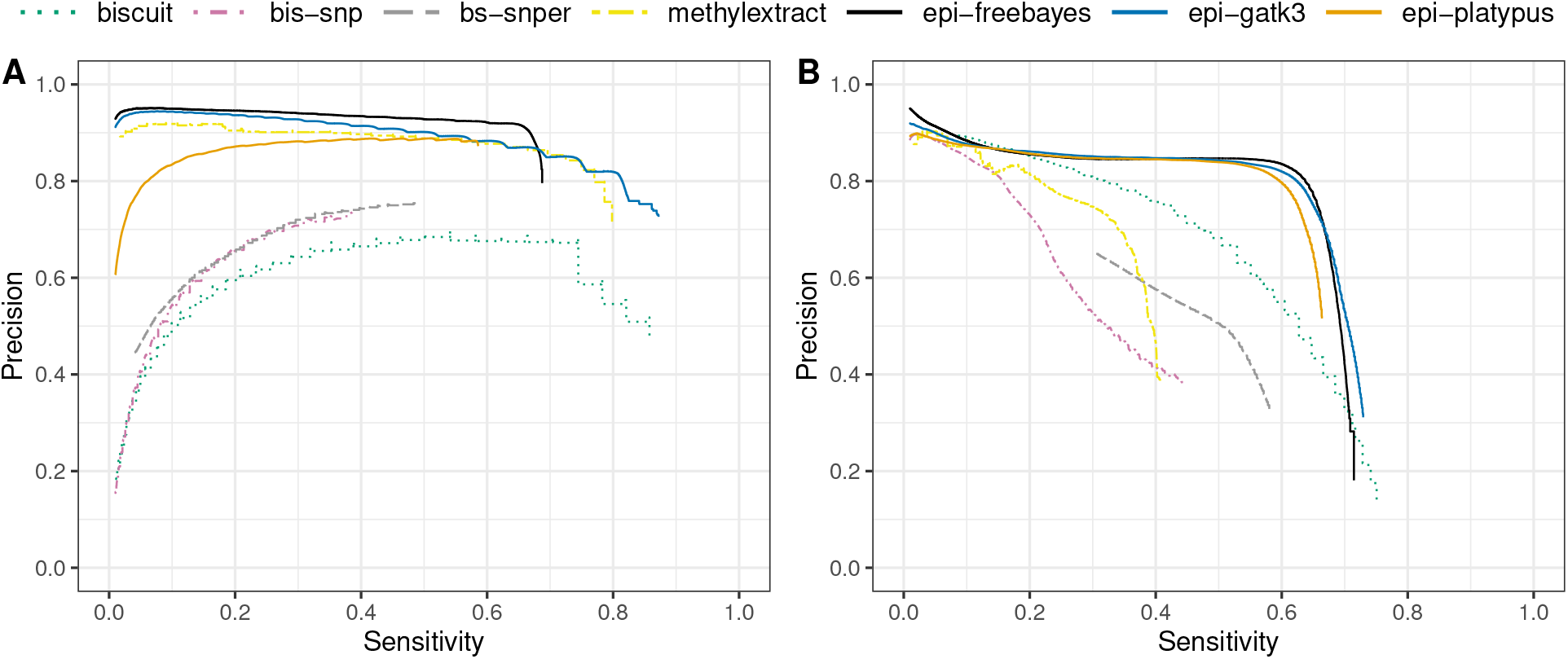
Precision-sensitivity of variants called in real data. In response to an increasing variant quality (QUAL) threshold, SNPs derived from published WGBS data are compared to those derived from established benchmark datasets for **(A)** *A. thaliana* and **(B)** human. Software with the epi-prefix are intended for conventional DNA sequencing libraries but in this case run after preprocessing with the double-masking procedure. True and false positives are evaluated based on both the substitution context and the estimated genotype.

**Figure 3:**
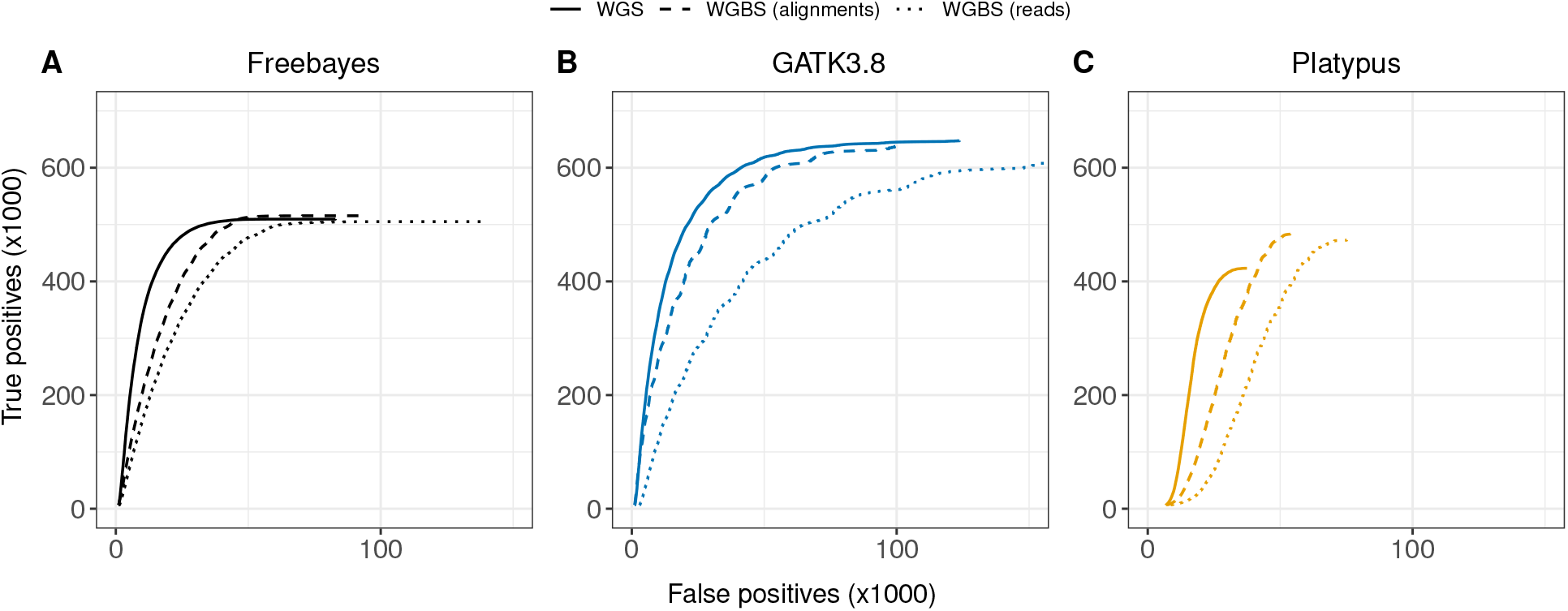
ROC-like comparisons in real and simulated data. In response to an increasing variant quality (QUAL) threshold, SNPs derived from real data (WGS) are compared to those derived from equivalent data (WGBS) after *in silico* bisulfite conversion of either reads or alignments, followed by preprocessing with the double-masking procedure, in *A. thaliana* (Cvi-0). Panels show results from conventional software **(A)** Freebayes, **(B)** GATK3.8 and **(C)** Platypus (default mode). True and false positives are evaluated based on both the substitution context and the estimated genotype.

The balance between precision and sensitivity can be evaluated using the harmonic mean to denote the F1 score, which can be compared between different software and data types (Table 1). With *in silico* WGBS reads, the optimal F1 scores for GATK3.8, Freebayes and Platypus were identified at 0.8508, 0.8039 and 0.7709, respectively, with a corresponding QUAL threshold of 80, 41 and 27. The overall best-performing tool was therefore GATK3.8, achieving 0.8685 sensitivity and 0.8338 precision at the optimal level, followed by Freebayes with 0.7335 sensitivity but a higher precision of 0.8894. Platypus performs better overall in default mode, despite an optimal precision level of 0.9436 for WGS and 0.8991 for WGBS data with assembly-mode enabled (not shown). The reduced overall performance due to lower sensitivity may in-part arise due to the need to set a pre-emptive threshold for Platypus at BQ≥0 (`--minBaseQual = 0`), following the double-masking procedure, to avoid over-filtering regions during local assembly.

**Table 1:** Optimised F1 scores in *A. thaliana* (Cvi-0). In comparison to the reference SNPs obtained from 1001 genomes, scores are derived when using real WGS and WGBS data, alongside *in silico* WGBS data derived from the WGS reads and alignments, respectively.

|  | Real data |  | *in silico* |  |
| --- | --- | --- | --- | --- |
|  | WGS | WGBS | reads | alignments |
| GATK3.8 | 0.9189 | 0.8177 | 0.8508 | 0.9069 |
| Freebayes | 0.8247 | 0.7670 | 0.8039 | 0.8247 |
| Platypus (default) | 0.7423 | 0.7026 | 0.7709 | 0.7935 |
| Platypus (assembly) | 0.6378 | 0.5980 | 0.6449 | 0.6509 |

The proportional deviation of false positives from *in silico* WGBS reads, relative to WGS data, show similar profiles when partitioned by substitution context (Supplementary Figure 2). A total of 92.3%, 77.3% and 72.8% of the total false positives here occur in positions which are homozygous-reference in the truth set for each of GATK3.8, Freebayes, and Platypus, respectively, after filtering those shared in the equivalent WGS data. These positions represent 12.0%, 5.6% and 5.6% of the total, unfiltered calls made by each tool. The remaining false positives typically comprise true variants which have been assigned an incorrect genotype (e.g. homozygous-alternative called as heterozygous), representing 2.9%, 4.2% and 4.6% of the total, unfiltered calls. Many of these cases suffer a low GQ likely as a consequence of reduced sequencing depth by limiting calls in bisulfite contexts to opposite-strand alignments. Such positions are also considered among the false negatives, alongside the fraction of true SNPs which are not called at all from bisulfite data. When considering the sequencing depth distribution of false negatives from *in silico* WGBS alignments, discounting those shared in the WGS data, there is a peak at ~4-5x in addition to a larger peak which correlates with the distribution for the true positives at ~18-20x (not shown). Accounting for a minimum per-position sequencing depth of ~7-10x should generally therefore be enough to make a successful call, disregarding differences due to WGBS alignment or significant deviations from typical sequencing biases (e.g. strand bias).

The indirect strand-specificity imposed on potential variant calls by the double-masking procedure is reflected by a reduction in the available sequencing depth required to make confident calls for potential polymorphisms involving thymine, in comparison to WGS data, which manifests predominately as a relative decrease in variant confidence metrics (i.e. QUAL and QualByDepth; described in Supplementary Table 1) on equivalent true positive SNPs (Figure 4). The number of true positive variants that would fail the recommended hard-filtering thresholds (QUAL<30 or QD<2.0), however, increased only from 1,730 (<0.27%) in WGS data to 9,762 (<1.55%) in the *in silico* WGBS data derived from WGS reads. There is also a minor increase in overall strand bias, as measured with the StrandOddsRatio (SOR) metric in GATK3.8 UnifiedGenotyper, where true positive variants that would fail the recommended hard-filtering threshold (SOR>3) increased from 18,045 (2.79%) in WGS data to 31,487 (5.0%) with simulated WGBS data. All together the number of true positive variants lost after hard-filtering increased from 30,858 (4.77%) to 56,695 (9.0%) due to *in silico* bisulfite conversion, while the total false positive variants increased from 80,528 (6.81%) to 143,745 (10.24%).

**Figure 4:**
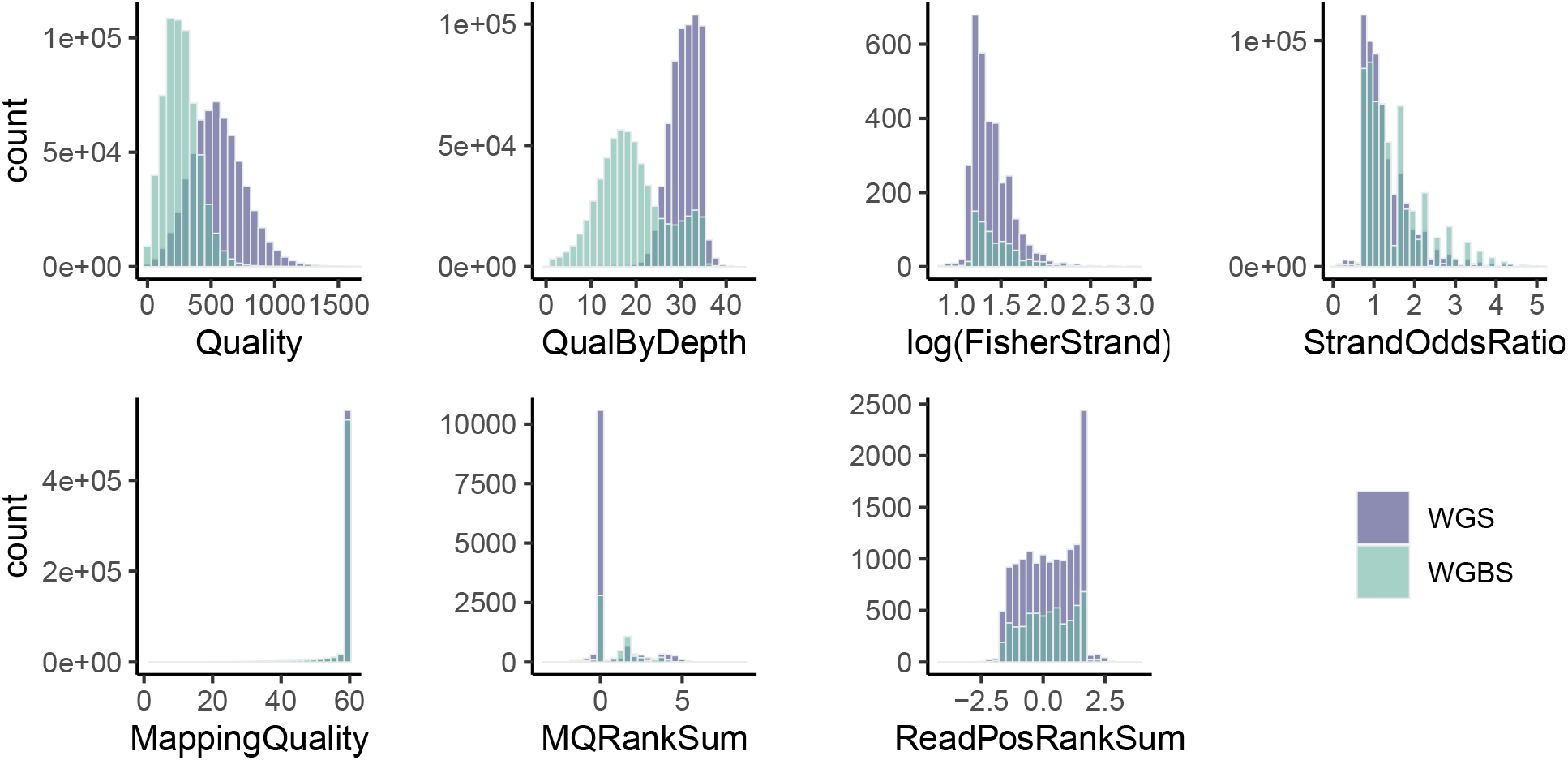
Quality control metric comparisons in real and simulated data. SNPs derived from real data (WGS) are compared to those derived from equivalent data (WGBS) after *in silico* bisulfite conversion of WGS reads, in *A. thaliana* (Cvi-0). Distributions are derived from the intersection of true positive calls made in each case by GATK3.8 UnifiedGenotyper, and each metric is considered specifically for hard-filtering in GATK best-practices. Definitions for each metric according to GATK are given in Supplementary Table 1. MQRankSum and ReadPosRankSum are only evaluated for a subset of calls, and in both datasets the vast majority of calls achieve a FisherStrand score of 0 (not shown).

## Discussion

Conventional germline variant callers can be broadly categorised as alignment-based, such as GATK3.8 UnifiedGenotyper, or haplotype-based, such as Freebayes and Platypus. Both strategies are concerned with correctly identifying variants at a given locus and inferring probabilistic genotype likelihoods based on allelic count differences, however they differ in their consideration of proximal variants to establish phase. Whilst UnifiedGenotyper considers precise alignment information in a position-specific, independent manner, Freebayes considers the literal sequence of each overlapping read to obtain the context of local phasing and derive longer haplotypes for genotyping. Some modern variant callers, including for example Platypus and GATK HaplotypeCaller, expand upon the haplotype-based approach by incorporating local assembly to aid in resolving potential indels. Bisulfite sequencing data can be made conceptually compatible with each of these described approaches, following pre-processing with the double-masking procedure, with the caveat that the chosen software for calling variants handles base quality specifically during the estimation of genotype likelihoods, ideally with an option for hard-filtering. Local assembly presents an added difficulty in that base quality is often considered additionally for read trimming during construction of De Bruijn graphs, e.g. in determination of “ActiveRegions” in GATK HaplotypeCaller, and is typically codependent on the same parameter used for setting its threshold during Bayesian inference. This can sometimes be circumvented, as demonstrated herein with Platypus, by allowing even a base quality of zero during local assembly before relying on the genotype likelihood model to weight such positions appropriately during variant calling, but such a case is not ideal. If masked nucleotides are allowed to be included in the model for deriving genotype likelihoods then the allelic balance on each variant will skew towards any mutations arising from bisulfite conversion, leading to a greater incidence of false positives.

It is important to consider that, unlike similar tools, variant calling with the presented approach is almost completely dissociated from the influence of cytosine methylation. The advantage of this is an improved sensitivity for high-confidence variants with fewer false positives, whilst preserving the underlying model of selected tools, but the methylation level itself must be evaluated independently. This is akin to several variant-independent approaches such as MethylDackel (https://github.com/dpryan79/MethylDackel) and GATK MethylationTypeCaller which are commonly used to estimate the methylation level without knowledge of the underlying SNPs. In combination with the presented approach it would be feasible to derive accurate variant-adjusted methylation calls, or even allele-specific methylation without the need for a corresponding genotype dataset obtained by conventional DNA sequencing.

In conclusion, the double-masking procedure facilitates sensitive and accurate variant calling directly from bisulfite sequencing data using software intended for conventional DNA sequencing libraries. The procedure can be readily adapted to existing software pipelines and does not necessitate any additional understanding of customised VCF files. Given sufficient sequencing depth, accurate alignment with minimal deviation from expected sequencing biases, and an appropriate level of filtering based on variant quality metrics, the SNPs derived from WGBS data are comparable to those from WGS data. The method presents a viable, alternative strategy to those who would otherwise need to sequence corresponding libraries of each type in order to better understand the role of DNA methylation in the context of the genetic background.

## Methods

### Validation datasets

All datasets analysed in this study are derived from published, public domain resources. High-quality reference variant datasets for human (NA12878) and *A. thaliana* (Cvi-0) accessions were obtained from Genome in a Bottle (GIAB) (https://ftp-trace.ncbi.nlm.nih.gov/giab/ftp/release/NA12878_HG001/NISTv4.2.1/) and the 1001 genomes project (https://1001genomes.org/data/GMI-MPI/releases/v3.1/), respectively. The corresponding reference genomes GRCh38 (GCF_000001405.26) and TAIR10 (GCF 000001735.3) were obtained from NCBI. Equivalent WGBS data were obtained from the NCBI Sequence Read Archive under accessions SRX3161707 and SRX248646. The original whole genome sequencing (WGS) data for *A. thaliana* Cvi-0 was also obtained, under accession SRX972441. Both trimmed reads and alignments from this accession were subject individually to *in silico* bisulfite treatment (~99% conversion rate) with custom in-house python scripts to generate corresponding, simulated WGBS datasets.

### Read processing and alignment

Reads were assessed with FastQC v0.11.8 (https://www.bioinformatics.babraham.ac.uk/projects/fastqc) and, where appropriate, trimming performed with cutadapt^23^ v2.5. WGS alignments were carried out with BWA^24^ v0.7.17-r1188, and WGBS alignments with BWA-meth^25^ v0.2.2. Read groups were merged with SAMtools^26^ v1.9, where appropriate, and PCR duplicates subsequently marked with Picard MarkDuplicates v2.21.1 (http://broadinstitute.github.io/picard).

### Variant calling

Following the double-masking procedure, variants were called using GATK^16^ v3.8 UnifiedGenotyper, Freebayes^17^ v1.3.1-dirty, and Platypus^18^ v0.8.1.2, in all cases with a hard filter of 1 on both minimum mapping quality (MAPQ) and BQ. Variants were called in addition using Platypus on assembly-mode with BQ≥0. For comparison, variants from the original bisulfite alignments were called also with BISCUIT v0.3.16.20200420 (https://github.com/huishenlab/biscuit), Bis-SNP^13^ v1.0.1, BS-SNPer^14^ v1.1 and MethylExtract^15^ v1.9.1. Default parameter settings were used, with the exception of minimum MAPQ and BQ thresholds which in all cases were set both to 1. The resulting variant calls were normalised, decomposed and otherwise processed for comparison to the high-quality reference data using BCFtools^26^ v1.9.

### Benchmarking

Benchmarking itself was performed with vcfeval of RTG Tools^27^ v3.11, which compares both the substitution context and estimated genotype of baseline variants from the truth set to each set of calls from bisulfite data in order to evaluate true positives, false positives and false negatives, in response to varying common filtering thresholds such as sequencing depth (DP), quality (QUAL) and genotype quality (GQ). Variants must occur with both the same substitution context and genotype in order to be evaluated as a true positive. Sensitivity refers to the true positives as a fraction of the truth set positives, whereas precision refers to the true positives as a fraction of the discovered variants (Supplementary Table 2). The F1 score reflects the balance of precision and sensitivity via the harmonic mean of both measures, and can be optimised relative to each filter by taking the maximum value in response to varying the relevant threshold.

## Data availability

All datasets used and/or analysed during the current study are derived from published, public domain resources. Accessions for reference genomes and sequencing data are specified in the methods, and corresponding datasets after *in silico* bisulfite conversion are available from the corresponding author on reasonable request. The human reference genome GRCh38 no alt plus hs38d1 analysis set and the benchmark VCF data can be obtained from the following URLs: https://ftp-trace.ncbi.nlm.nih.gov/giab/ftp/release/references/GRCh38/; https://ftp-trace.ncbi.nlm.nih.gov/giab/ftp/release/NA12878_HG001/NISTv4.2.1/; https://1001genomes.org/data/GMI-MPI/releases/v3.1/

## Code availability

The described method is available at https://github.com/bio15anu/revelio. An example implementation of the method is also provided at https://github.com/EpiDiverse/SNP, together with installation instructions, documentation and a demo dataset. All software is open-source and licensed under the MIT license.

## Acknowledgements

We would like to thank all the members of the EpiDiverse Consortium for their active and invaluable support in discussing, developing and performing these analyses. Special thanks to Morgane Van Antro and Samar Fatma for their assistance in optimising the benchmarking procedure. The European Training Network “EpiDiverse” received funding from the EU Horizon 2020 program under Marie Skłodowska-Curie grant agreement No 764965.

## Author contributions

CO, PFS and DL identified the gap to be addressed by a new method. All authors were involved in discussions. AN and CO devised the method and the following benchmark analysis. AN and MF implemented a working pipeline implementation of the method. AN performed the benchmarking analysis, and interpreted results with CO. AN wrote, and all authors read and approved the final manuscript.

## Supporting information

Supplementary Tables 1-2 and Supplementary Figures 1-2 available online.

## Algorithm 1 The double-masking algorithm performed on each alignment.

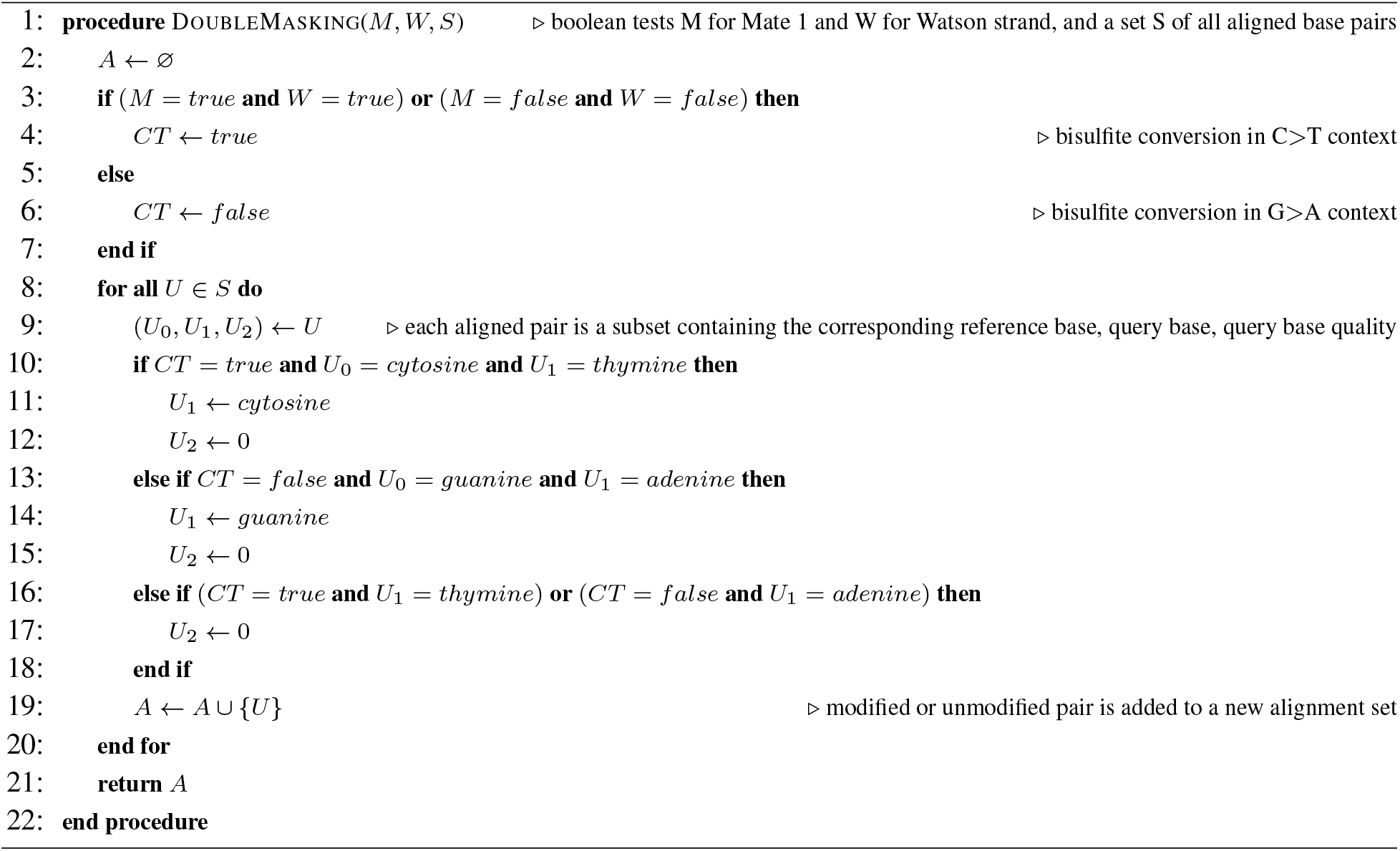

