## Supplementary Material for "Manipulating base quality scores enables variant calling from bisulfite sequencing alignments using conventional Bayesian approaches"

| **Supplementary Table 1. Summary of QC metrics as described by GATK.** Definitions obtained from <https://gatk.broadinstitute.org/hc/en-us/articles/360035890471> (accessed 8th January 2020). Filtering thresholds are according to best-practices. | | |
| --- | --- | --- |
| Metric | Threshold | Summary |
| **Quality**  **(QUAL)** | < 30 | Variant confidence. The Phred-scaled probability that there is some kind of nucleotide variation at a given site. |
| **QualByDepth**  **(QD)** | < 2 | The variant confidence (from the QUAL field) divided by the unfiltered depth of non-hom-ref samples. Intended to normalize the variant quality in order to avoid inflation caused when there is deep coverage. |
| **FisherStrand**  **(FS)** | > 60 | The Phred-scaled probability that there is strand bias at the site, indicating whether the alternate allele was seen more or less often on the forward or reverse strand than the reference allele. |
| **StrandOddsRatio**  **(SOR)** | > 3 | Another way to estimate strand bias using a test similar to the symmetric odds ratio test. Created because FS tends to penalize variants that occur at the ends of exons. |
| **RMSMappingQuality**  **(MQ)** | < 40 | The root mean square mapping quality over all the reads at a given site. |
| **MappingQualityRankSumTest (MQRankSum)** | < -12.5 | The u-based z-approximation from the Rank Sum Test for mapping qualities. It compares the mapping qualities of the reads supporting the reference allele and the alternate allele. |
| **ReadPosRankSumTest**  **(ReadPosRankSum)** | < -8 | The u-based z-approximation from the Rank Sum Test for site position within reads. It compares whether the positions of the reference and alternate alleles are different within the reads. |

| **Supplementary Table 2. The confusion matrix.** The table shows the relationship between true and false positives. Precision is calculated by taking the true positives as a proportion of the predicted condition positives, and sensitivity is calculated by taking the true positives as a proportion of the true condition positives. The F1 score is the harmonic mean of precision and sensitivity. Equations denoted in the footnote. | | | |
| --- | --- | --- | --- |
|  |  | **True Condition** | |
|  |  | Positives | Negatives |
| **Predicted Condition** | Positives | True Positives | False Positives  (Type II error) |
|  | Negatives | False Negatives  (Type I error) | True Negatives |

$$precision= {True positives}/\left( True positives+False positives \right)$$

$$sensitivity= {True positives}/\left( True positives+False negatives \right)$$

$$F1 score= 2\cdot\left( \left( recall\cdot precision \right)/\left( recall+precision \right) \right)$$

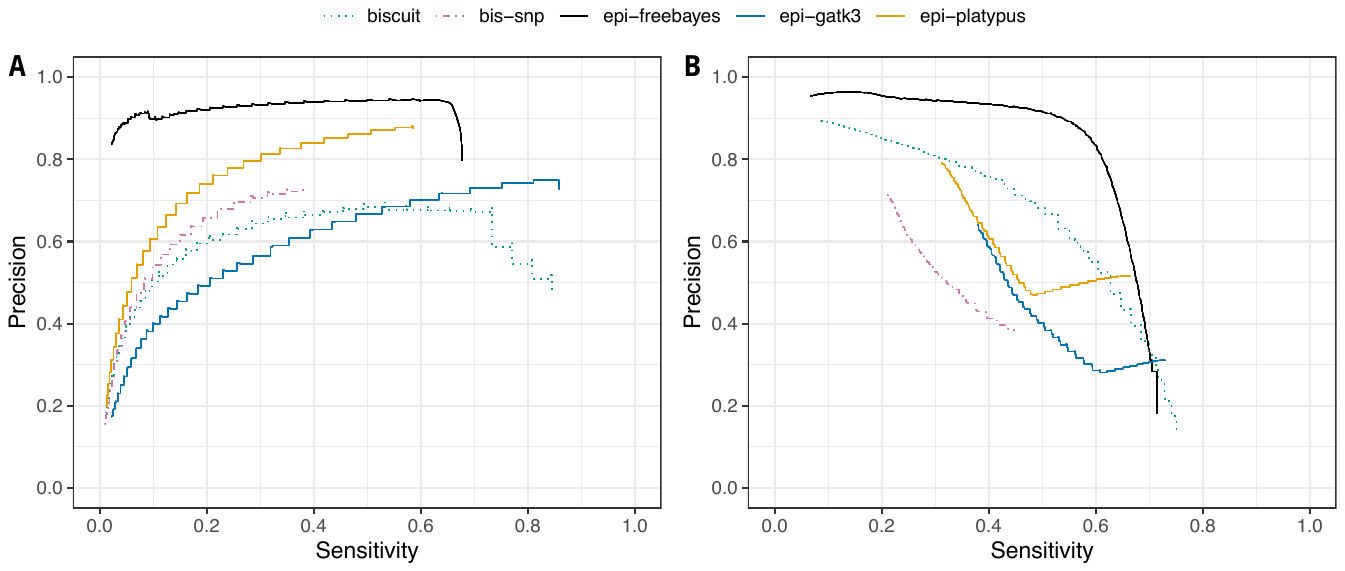

**Supplementary Figure 1. Precision-sensitivity of variants called in real data.** SNPs derived from published WGBS data are compared to those derived from established benchmark datasets for both **(A)** *A. thaliana* and **(B)** human. True and false positives are evaluated based on both the substitution context and the estimated genotype. Software with the epi- prefix are intended for standard sequencing data but in this case run with alignments pre-processed with the double-masking procedure. With genotype quality (GQ) as a qualifier, epi-freebayes performs consistently high in terms of the optimal balance of true and false positives, with an F1 score of 0.7715 and 0.6984 in *A. thaliana* and human datasets, respectively. BS-SNPer and MethylExtract do not give values for genotype likelihood or genotype quality in the output VCF files. Note that the given GQ values in biscuit are identical to the QUAL values in single-sample mode.

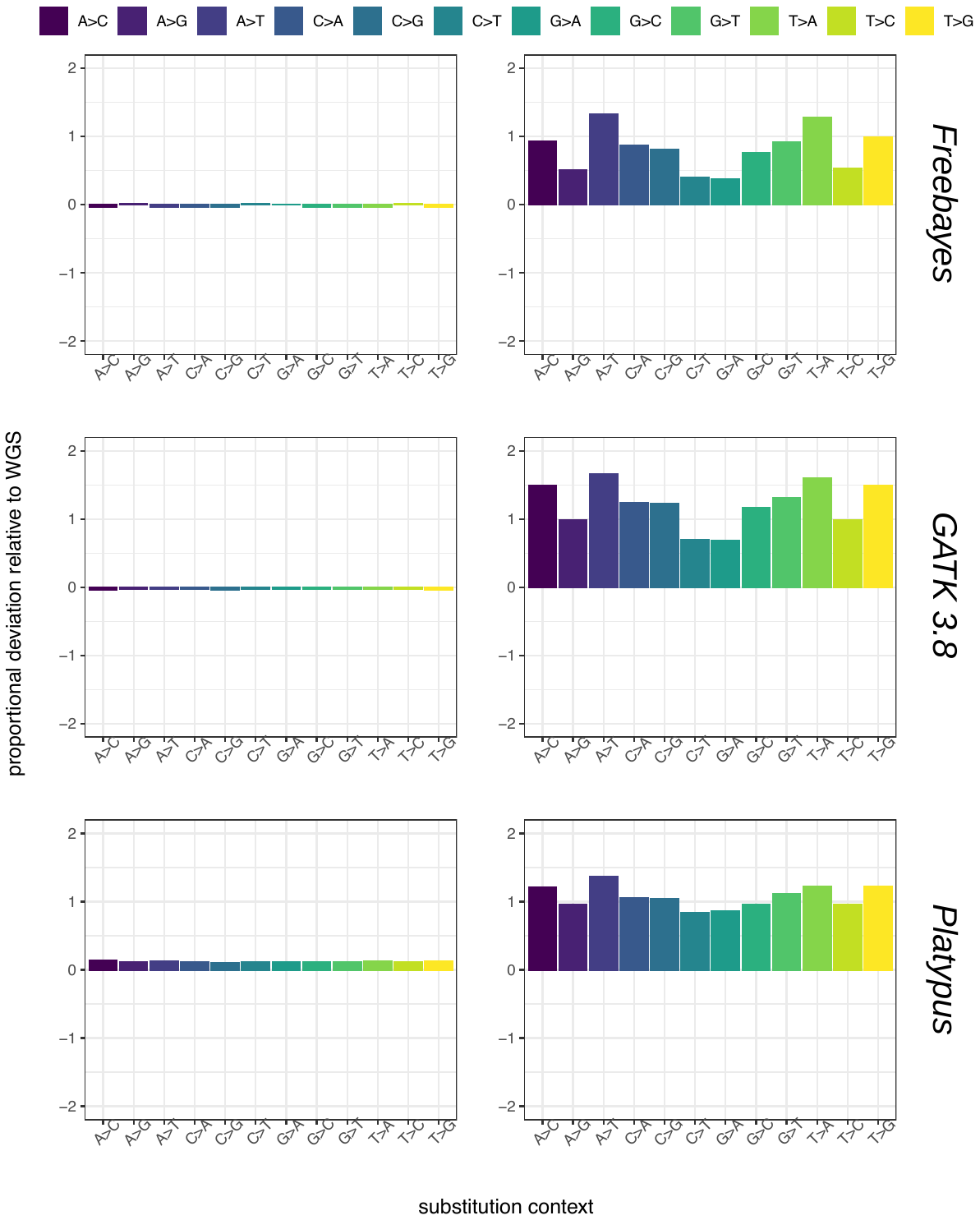

**Supplementary Figure S2. Proportional deviation of variant contexts in real and simulated data.** Raw, unfiltered variant calls from data subject to *in silico* bisulfite treatment (WGBS) are shown relative to the original untreated WGS data, in *A. thaliana* (Cvi-0). The left-hand panels indicate the fraction of true positives in each dataset, whereas the right-hand panels refer to the fraction of false positives. The profiles partitioned across each substitution context are similar between each tool, although there are fewer false positives in Platypus (default mode) and Freebayes relative to GATK 3.8, and in contrast to the other tools the true positives called by Platypus seem to increase overall in the artificial WGBS data relative to WGS.
